# Remodeling of the protein ubiquitylation landscape in the aging vertebrate brain

**DOI:** 10.1101/2023.12.02.569713

**Authors:** Antonio Marino, Domenico Di Fraia, Diana Panfilova, Amit Kumar Sahu, Alessandro Ori

**Author notes:** Equal contribution (Antonio Marino, Domenico Di Fraia).

## Abstract

Post-translational modifications (PTMs) regulate protein homeostasis and function. How aging affects the landscape of PTMs remains largely elusive. Here, we reveal changes in hundreds of protein ubiquitylation, acetylation, and phosphorylation sites in the aging brain of mice. We show that aging has a major impact on protein ubiquitylation and that 29% of the ubiquitylation sites are affected independently of protein abundance, indicating altered PTM stoichiometry. We found a subset of these sites to be also affected in the brain of the short-lived killifish *Nothobranchius furzeri*, highlighting a conserved aging phenotype. Furthermore, we estimated that over 35% of ubiquitylation changes observed in old mouse brains derive from partial proteasome inhibition, a well-established hallmark of brain aging. Our findings provide evidence of an evolutionarily conserved ubiquitylation signature of the aging brain and establish a causal link between proteasome inhibition and age-related remodeling of the ubiquitylome.

## Introduction

Posttranslational modifications (PTMs) expand the chemical diversity of proteins by generating distinct proteoforms encoded by the same gene (L. M. Smith et al., 2013). PTMs modulate the function of proteins by regulating their localization, interactions, stability, and turnover, thereby influencing protein homeostasis (proteostasis) (Zhong et al., 2023). Loss of proteostasis is a hallmark of aging (Santra et al., 2019; C. B. Smith et al., 1995; Taylor et al., 2011) and age-related diseases, particularly neurodegenerative disorders (Hou et al., 2019; Vilchez et al., 2014), that ultimately manifests with the formation of protein aggregates. Acetylation (Ac), phosphorylation (Phospho), and ubiquitylation (Ub) are among the most studied PTMs and account for more than 90% of all the reported modifications in the dbPTM database (Ramazi et al., 2021). Acetylation is fundamental for brain gene expression, regulating the accessibility of histones (Toker et al., 2021). Histone acetylation is essential for neuronal maturation, synapse formation, and establishment of neuronal circuits (Bonnaud et al., 2016). Phosphorylation mediates signaling and takes part in virtually every neuronal process. For instance, calcium/calmodulin-dependent protein kinase II (CaMKII) modulates synaptic strength, mainly by affecting the trafficking, functional properties, and anchoring to the postsynaptic membrane of glutamate transmembrane receptors (Lisman et al., 2002). Ubiquitylation plays a central role in protein degradation through the ubiquitin-proteasome system (UPS) and autophagy/lysosome pathway (R.-H. Chen et al., 2019; Kirkin et al., 2009; Tai et al., 2008), two proteostasis components whose activity declines with aging (Kaushik et al., 2012; Kelmer Sacramento et al., 2020; Yu et al., 2017). Ubiquitin signaling is also required for the turnover of organelles, e.g., mitochondria via mitophagy and endoplasmic reticulum (ER) via ER-phagy (González et al., 2023; Pickles et al., 2018), and it can modulate synaptic activity and plasticity (Hallengren et al., 2013; Mabb et al., 2010).

Proteins carrying PTMs, including acetylation, phosphorylation, and ubiquitylation, were found in protein aggregates in samples from patients suffering from different types of neurodegenerative diseases (Kabir et al., 2022; Perluigi et al., 2016; Schmidt et al., 2021). For instance, MAPT (Tau) hyperphosphorylation and ubiquitylation are characteristic of Alzheimer’s (AD) (Lowe et al., 1988; Perluigi et al., 2016). TDP-43 was found to be ubiquitylated in frontotemporal dementia (FTD) (Arai et al., 2006; Neumann et al., 2006), while GFAP acetylation has been reported for amyotrophic lateral sclerosis (ALS) (Liu et al., 2013).

A few previous studies investigated proteome-wide changes in PTMs during vertebrate brain aging. Cysteine oxidation in mice was analyzed by (Xiao et al., 2020), showing the presence of age– and tissue-specific redox-regulated clusters, e.g., tRNA aminoacylation complex, and found no relationship between changes in protein oxidation and abundance with age. (Petrovic et al., 2021) reported that protein persulfidation, another redox modification, decreases during rat brain aging and neurodegenerative disorders. Additionally, in rats, phosphorylation data showed that specific phosphosite levels changed in the aging brain due to mis-localized protein kinases (Ori et al., 2015). These studies provided evidence that altering PTMs might contribute to the loss of protein homeostasis in aging and age-related neurodegenerative disorders. However, a systematic investigation of major PTMs during physiological brain aging is still lacking.

To fill this knowledge gap, we quantified the effect of aging on protein acetylation, phosphorylation, and ubiquitylation in the mouse brain using mass spectrometry. We found ubiquitylation to be the most affected PTM. Given the conservation of ubiquitin across evolution (Zuin et al., 2014), we asked whether the same alterations could be observed in different species. Thus, we turned to the short-lived *Nothobranchius furzeri* (killifish) because of its pronounced age-related neurodegeneration phenotypes, including accumulation of phosphorylated Tau with aging (de Bakker et al., 2023; Di Fraia et al., 2023; Louka et al., 2022). Notably, both mice and killifish exhibit an age-related accumulation of ubiquitin (Matsui et al., 2019; Ohtsuka et al., 1995) and a decline in proteasome activity (Burov et al., 2023; Kelmer Sacramento et al., 2020). By combining mouse and killifish data, we identified a ubiquitylation aging signature conserved in the brains of these two species. Finally, we used human induced pluripotent stem cell (iPSC)-derived neurons (iNeurons) to demonstrate that a significant portion of the observed age-related changes of ubiquitylation can be attributed to a decline in proteasome activity.

## Aging affects PTMs in the brain

To assess the impact of aging on PTMs in the mouse brain, we used a workflow based on label-free Data Independent Acquisition (DIA) mass spectrometry to analyze from the same set of samples three major modifications: ubiquitylation, phosphorylation, and acetylation (Figure 1A, see methods). For ubiquitylated peptides enrichment, we used lysine di-GLY (K-ε-GG) remnant motif pulldown (Fulzele et al., 2018). This method leads to the enrichment of other modifications like NEDDylation and ISGylation. However, more than 95% of K-ε-GG-modified sites have been shown to come from ubiquitin (Kim et al., 2011). From this point forward, we refer to K-ε-GG-modified sites as ubiquitylated. We quantified changes for 10487, 7031, and 6049 phosphorylation, ubiquitylation, and acetylation sites, respectively. In parallel, we measured differences in total protein and transcript abundance, covering 6453 proteins and 17994 transcripts. Our datasets recapitulated changes in PTMs that have been previously associated with aging and disease, e.g., MAPT (Tau) hyperphosphorylation and GFAP acetylation (Gong et al., 2008; Liu et al., 2013) (Figure S1 and Table S1).

**Figure 1.**
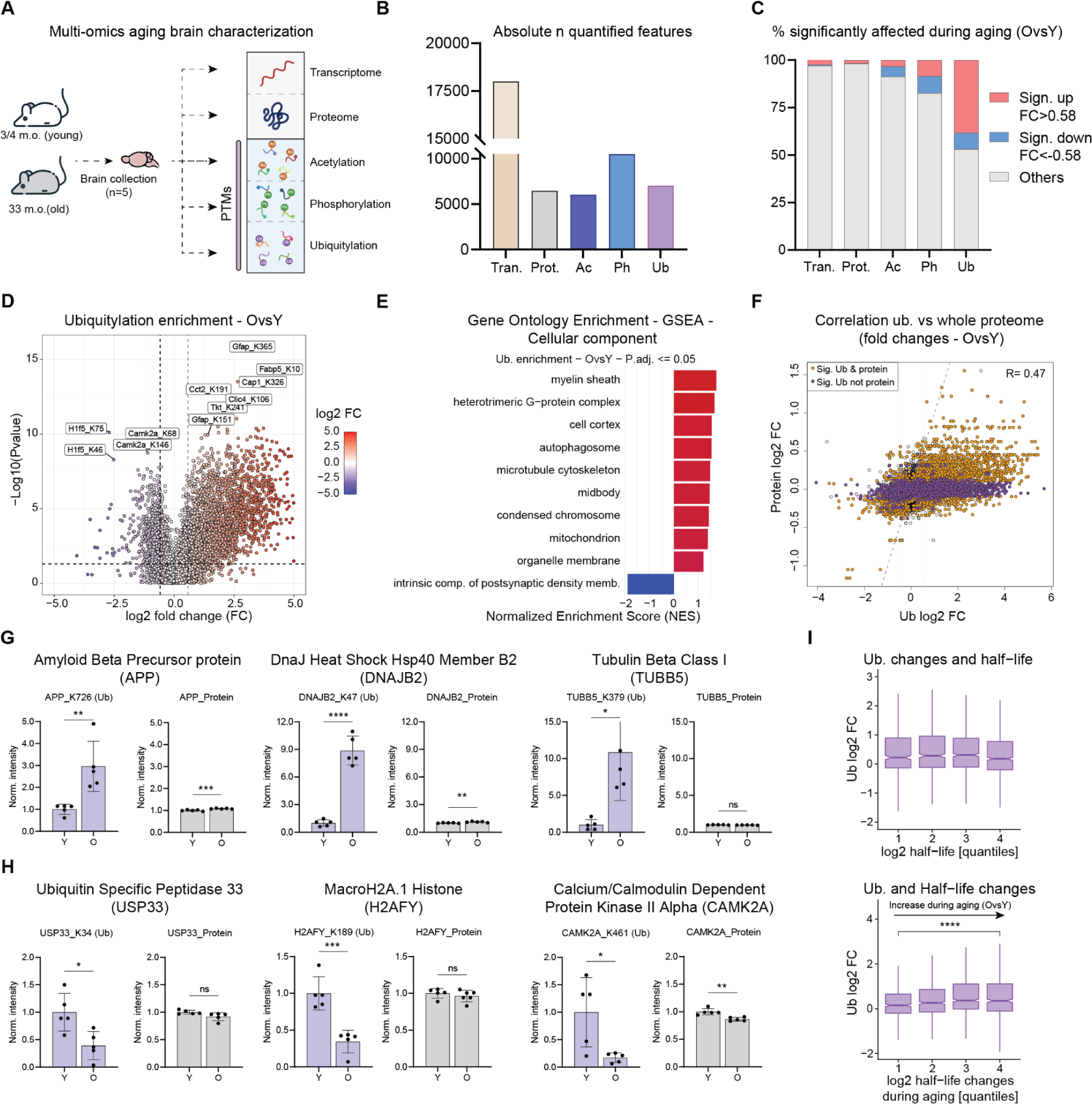
PTM landscape of the mouse aging brain. A) Multi-omics approach scheme used to characterize mouse brain aging. B) Absolute number of quantified features in each omic layer. C) Percentage of significantly affected transcripts, proteins, or PTMs (Adj.PValue<0.05 for proteome and transcriptome, QValue<0.05 for PTMs; fold change cutoff: FC>0.58, FC<-0.58). D) Volcano plot for ubiquitin enrichment in mouse brain aging (N=5). E) GO enrichment analysis for ubiquitin enrichment using GSEA Biological processes (Adj.PValue<0.05). F) Scatterplot describing the correlation between proteome and ubiquitylome, in orange are highlighted proteins which are affected significantly in both layers (Ub QValue<0.05; proteome Adj.PValue<0.05), in purple proteins which are significantly changing only for ubiquitylation (Ub QValue<0.01; proteome Adj.PValue>0.05). G) and H) Examples of proteins with an increase or decrease level ubiquitylation independently of protein abundance (N=5, unpaired *t*-test). I) Correlation between ubiquitylation changes and protein half-life (Fornasiero et al., 2018) (upper panel) or changes in protein half-life (Kluever et al., 2022) during mouse brain aging (bottom panel) (Wilcoxon test). *P ≤ 0.05; **P ≤ 0.01, ***P ≤ 0.001, ****P ≤ 0.0001. Related to Figure S1, S2, S3 and Table S1.

Among the analyzed PTMs, we observed a striking effect on the percentage of ubiquitylation sites significantly affected by aging (Figure 1B-C). The ubiquitin dataset’s principal component analysis (PCA) consistently depicted a clear separation difference between young and old age groups (Figure S2A). A higher number of ubiquitylation sites were identified in old samples (Figure S2B), and age-related changes of ubiquitylation were skewed towards positive values (Figure 1D). These observations are consistent with previous observations describing increased high-molecular-weight ubiquitylated conjugates but not free ubiquitin in mouse brain (Gray et al., 2003; Ohtsuka et al., 1995), which we further validated in our samples via dot-blot analysis (Figure S2C). Despite the overall trend of increased ubiquitylation with aging, a fraction of sites showed decreased ubiquitylation (Figure 1E). When assessing which cellular component categories were affected by these changes, GO enrichment analysis revealed that the myelin sheath, mitochondrion, and GTPase complex showed an increase in ubiquitylation, the latter in agreement with a previous report (Lei et al., 2021) (Figure 1E). In contrast, proteins belonging to the synaptic compartment were enriched among those showing decreased ubiquitylation with aging (Figure 1E). We did not find these changes reflected by proteome and transcriptome datasets that, instead, highlighted a prominent inflammation signature (Figure S3), a well-described hallmark of aging (Yin et al., 2016).

Next, we asked whether changes in ubiquitylated peptide abundance could result from underlying alterations of total protein abundance. Although changes in ubiquitylation were positively correlated with changes in protein abundance (R=0.47; pValue<2.2e-16), 29% of the altered sites could not be explained by altered protein abundance. This observation indicates that changes in ubiquitylation site occupancy occur in the aging brain (Figure 1F). Among cases that showed a prominent increase in ubiquitylation independently of protein levels, we found proteins encoded by neurodegeneration-associated genes, e.g., APP, TUBB5, and chaperones/ co-chaperones, e.g., DNAJB2 (Figure 1G). In contrast, several histones, deubiquitinases, and synaptic proteins, including H2AFY, USP33, and CAMK2A, showed reduced ubiquitylation (Figure 1H). Interestingly, we also identified multiple proteins that contained ubiquitylated sites affected in the opposite manner (Figure S2D-G), indicating a complex rewiring of protein ubiquitylation in the aging mouse brain.

Since ubiquitin-mediated degradation can regulate protein half-life, we correlated our protein ubiquitylation data with protein half-life (Fornasiero et al., 2018) and protein half-life changes during aging measured in mouse brain (Kluever et al., 2022). Interestingly, an increase in ubiquitylation correlated with an increase in protein half-life with aging but not with half-life itself (Figure 1I), suggesting that the accumulation of ubiquitylated proteins might reflect an age-related impairment in the protein clearance capacity of the brain. Together, these results show ubiquitylation is one of the most prominent proteome differences between young and old brains. Thousands of ubiquitylation sites undergo changes in site occupancy independently from protein level alterations. Additionally, our results indicate that decreased protein turnover might be associated with an accumulation of ubiquitylated proteoforms in the aging brain.

## Conserved ubiquitylation changes in the aging vertebrate brain

Next, we investigated whether age-related alterations in ubiquitylation are conserved across different species by comparing our mouse dataset with one we previously generated for the short-lived killifish *Nothobranchius furzeri* (Di Fraia et al., 2023). We hypothesized that any shared changes could reveal a fundamental ubiquitylation signature universally linked to vertebrate brain aging. First, we confirmed that also in killifish, differences in ubiquitylation are largely independent of protein abundance (Figure S4A). Next, we aligned mouse and killifish ubiquitylated sites (see methods and Table S2). We compared age-related ubiquitylation changes in the two species upon correction for any underlying total protein differences that might confound the comparison (as described in (Di Fraia et al., 2023)). Of the 1640 sites we could align between mouse and killifish, 58% changed consistently or showed a consistent trend when relaxing filtering criteria (p<0.25 in both datasets, Figure 2A).

**Figure 2.**
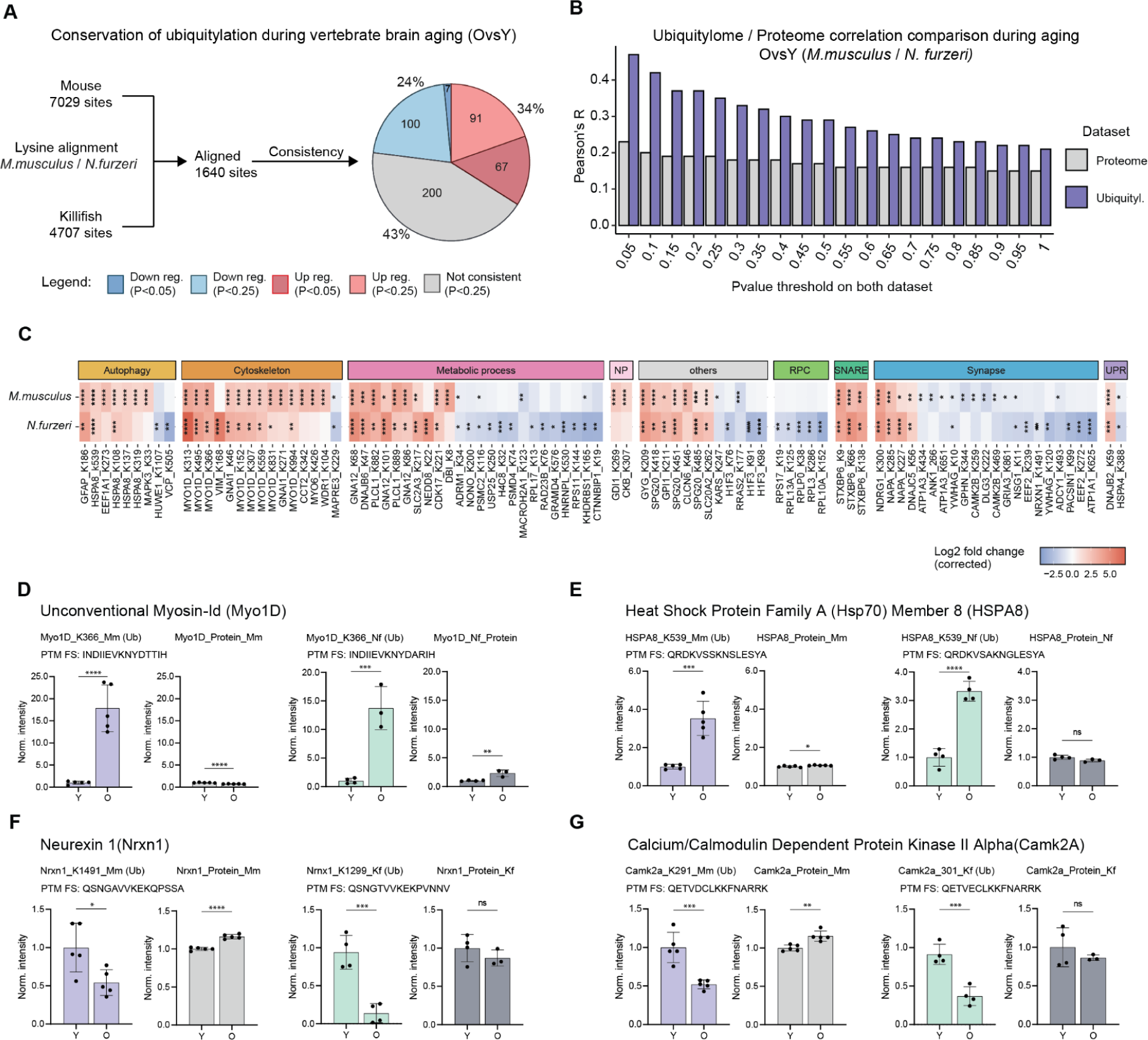
Conservation of age-related ubiquitylation changes in killifish and mouse. A) Mapping of ubiquitylation sites between *M. musculus* and *N. furzeri*. The percentage of consistently up (red and light red for Pvalue<0.05 or PValue<0.25, respectively, in both datasets), down (blue and light blue for Pvalue<0.05 or PValue<0.25, respectively, on both datasets), and not consistently (gray; PValue<0.25) regulated sites are shown in the pie plot. B) Ubiquitylome and proteome correlation between mouse and killifish. Purple bar plots show the correlation between the two species’ ubiquitylomes, and gray bar plots instead the correlation between the proteomes. On the y-axis, different PValue thresholds are applied. C) Heatmap of sequence-aligned peptides ubiquitylation for different cellular compartments in mouse and killifish. Fold changes have been corrected for protein abundance (PValue<0.05 in at least one species). D-G) Examples of peptides with conserved increased or decreased levels of ubiquitylation independently of protein abundance, flanking sequence (FS) of the modified peptides was added under the protein name (mouse N=5, killifish N=3-4, unpaired *t*-test). *P ≤ 0.05; **P ≤ 0.01, ***P ≤ 0.001, ****P ≤ 0.0001. Related to Figure S4 and Table S2.

Additionally, we examined the correlation between age-related ubiquitylation and proteome differences to determine their conservation across the two species. We found a higher concordance between ubiquitylation changes (expressed as Pearson’s correlation between log2 fold changes) than their respective proteomes, independently of the significance cut-off used (Figure 2B, S4B-C). When we analyzed which categories of proteins were consistently affected, we identified a subset of proteins related to synapse, autophagy, cytoskeleton, and metabolism (Figure 2C).

Furthermore, we observed that some of these proteins showed more than one conserved affected site, e.g., MYO1D, HSPA8, and CAMK2A (Figure 2D-G). MYO1D has been involved in autophagy regulation (Feng et al., 2022), while HSPA8’s primary function is to facilitate protein folding but has been reported to prevent tau fibril elongation (Baughman et al., 2018). CAMK2A, which decreases in ubiquitylation, is an essential synaptic plasticity regulator (Lisman et al., 2002). The robust and conserved effects observed for these proteins suggest them as potential ubiquitylation biomarkers of brain aging.

## Proteasome inhibition can partially recapitulate age-related ubiquitylation changes

Having defined a conserved remodeling of protein ubiquitylation landscape in the aging brain, we aimed to identify its molecular causes. Since reduced proteasome activity is an established hallmark of brain aging observed in mammals and other species (Burov et al., 2023; Gray et al., 2003; Hipp et al., 2019; Kelmer Sacramento et al., 2020), we wanted to assess its contribution to the perturbation of protein ubiquitylation in a relevant model system. We decided to use human iPSC-derived neurons (iNeurons) (Wang et al., 2017) because they are been established as a reference *in vitro* model for human age-associated neurodegenerative disorders (Pantazis et al., 2022), and have previously been used for proteome-wide investigations of protein ubiquitylation (Antico et al., 2021; Ordureau et al., 2020). Thus, we impaired proteasome activity in 7 days and 14 days post-differentiation in iNeurons using bortezomib for 24 hours. To assess in parallel the contribution of the lysosome/autophagy pathway, we also analyzed iNeurons treated with the lysosomal V-ATPase inhibitor bafilomycin, which causes reduced acidification of lysosomes and consequent inhibition of lysosomal proteases, and a third group treated by both compounds (Figure 3A). No overt cellular toxicity was observed after 24 hours of treatment. However, we noted thinner neuronal projections in bafilomycin-treated neurons (Figure S5A). We validated the treatments using immunoblot, showing the expected increase of K48-linked ubiquitin chains and SQSTM1 (p62) protein levels upon proteasome inhibition and increased LC3B-II/LC3B-I ratio following bafilomycin treatment (Figure S5B). Next, we generated mass spectrometry data to quantify changes in protein abundance and K-ε-GG-enriched peptides. Principal component analysis of proteome data highlighted differentiation day as the most prominent signature, although the effect of the different treatments could be observed (Figure 3B).

**Figure 3.**
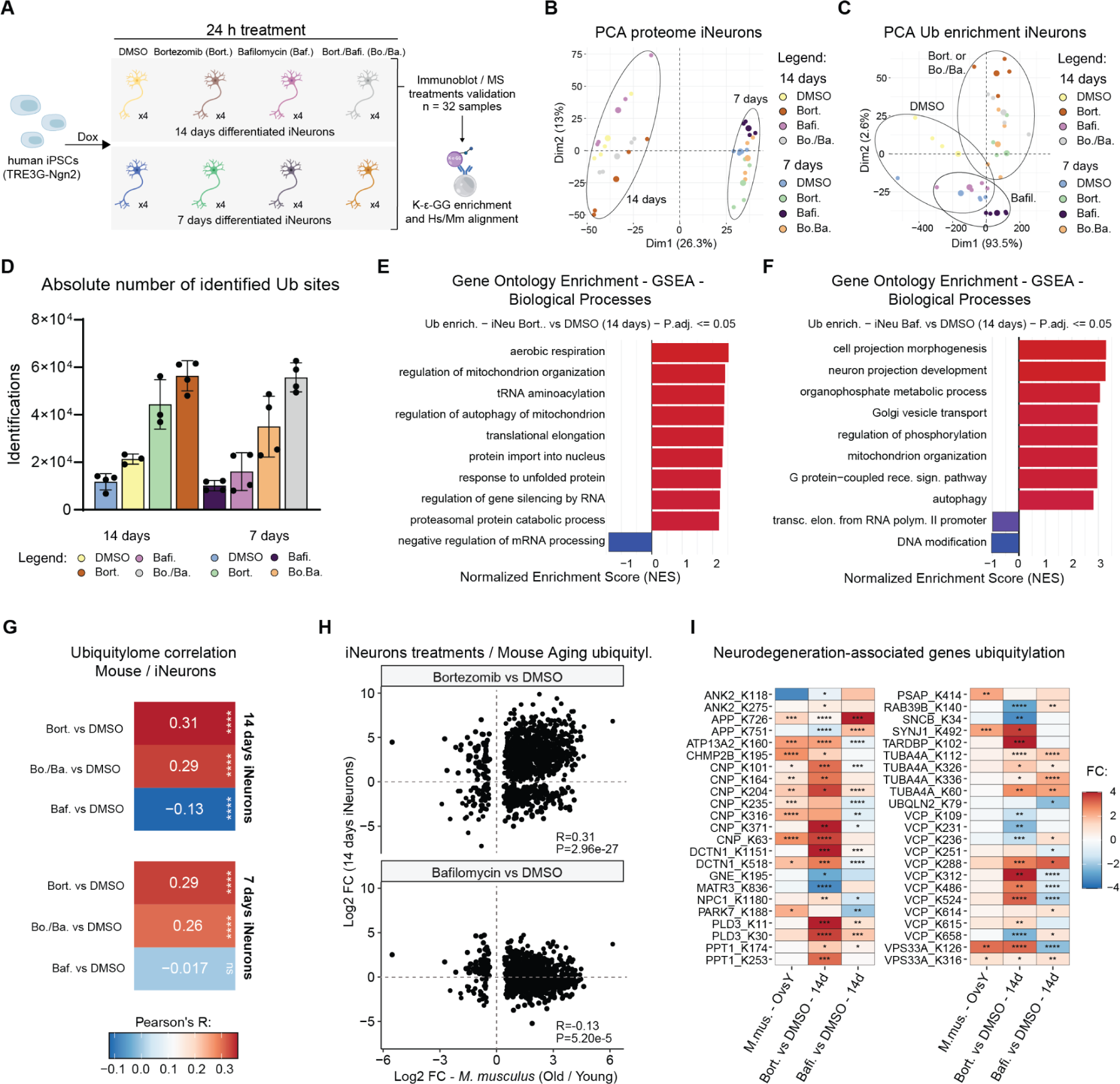
Proteasome inhibition leads to neurodegeneration-associated protein ubiquitylation in iNeurons. A) Experimental scheme for iNeurons drug treatment and characterization (N=4 for each condition obtained from two different differentiation batches). B) Principal component analysis (PCA) of proteome changes in 7 and 14 days iNeurons, the ellipses highlight the differentiation days groups. C) PCA of ubiquitylation changes in 7 and 14 days iNeurons, the ellipses highlight the differentiation drug treatments. D) Barplots showing the absolute number of identified ubiquitylated sites in the different sample groups. E-F) GO enrichment analysis for ubiquitin enrichment using GSEA Biological processes, on the left 14 days iNeurons treated with bortezomib compared to DMSO on the right 14 days iNeurons treated with bafilomycin compared to DMSO (P.adj.Value<0.05). G) Correlation between mouse aging and 14 days iNeurons bortezomib and bafilomycin datasets (PValue<0.05 on both datasets; Pearson correlation test). H) Scatterplot for ubiquitylome comparison between mouse aging and 14 days iNeurons treated with bortezomib (upper panel) or bafilomycin (lower panel) compared to DMSO (PValue<0.05 on both datasets). I) Heatmap of peptides changes in the ubiquitylation of neurodegeneration-associated genes significant (PValue<0.05) in at least one condition; the lysine numbering refers to the mouse protein sequence. *P ≤ 0.05; **P ≤ 0.01, ***P ≤ 0.001, ****P ≤ 0.0001. Related to Figure S5, S6, S7 and Table S3 and S4.

Conversely, data from K-ε-GG-enriched peptides highlighted a more substantial impact of drug treatments over differentiation days (Figure 3C). As expected, we noted a robust increase in the number of identified ubiquitylated peptides upon bortezomib treatment, while the number of total proteins remained similar (Figure 3D, S5C, and Table S3). GO enrichment analysis performed using proteins that displayed changes in ubiquitylation showed that the drug treatments affected different cellular compartments. Bortezomib increased the modification of the mitochondrial and unfolded protein response-related proteins and caused a decrease in the ubiquitylation of factors involved in mRNA maturation (Figure 3E). Instead, bafilomycin treatment predominantly affected proteins related to neuron projection and Golgi vesicle transport (Figure 3F). When we compared ubiquitylation and protein abundance changes, we noted a lower correlation than the one we observed in brain tissue (Figure S6). This difference might be due to the acute treatment modality performed in iNeurons, which renders the contribution of protein abundance changes to ubiquitylation negligible (Figure S7). Hence, we did not apply any additional correction for this particular dataset. To better understand the correlation between the signatures induced by the two drug treatments on cultured neurons and the protein modifications observed *in vivo* during aging, we aligned human lysine residues to the corresponding mouse and killifish orthologs. We could map 57% of the ubiquitylated lysines for mouse (3961 sites) and 44% for killifish (2053 sites) across all iNeurons treatment conditions (Table S4).

When we correlated iNeurons and mouse aging significantly (P<0.05) affected sites, we observed that proteasome inhibition, but not lysosomal acidification impairment, could replicate a substantial portion of the ubiquitylation aging signature (Figure 3G). The majority of the age-related changes in ubiquitylation in the mouse dataset (∼60% of the affected sites, P<0.05) showed congruent changes (P<0.05) in response to the proteasome inhibition treatment (iNeurons, 14 Days). Interestingly, among the shared alterations, SQSTM1 (p62), which is involved in the formation and autophagic degradation of cytoplasmic ubiquitin-containing inclusions (Clausen et al., 2010), showed a prominent increase in ubiquitylation in mouse brain aging and, as expected, in the bortezomib-treated 14 days iNeurons (Figure S5E). Although to a lower extent, the same trends were observed for killifish both in terms of global correlation (Figure S5E-F) and percentage of affected sites (∼42%, P<0.05 against iNeurons, 14 Days). These results show that partial proteasome inhibition can recapitulate a significant portion of the ubiquitylation changes occurring *in vivo* during aging, showing a higher correlation and percentage of ubiquitylated sites consistently regulated compared to lysosomal acidification impairment (Figure 3H).

Finally, as ubiquitylation is a marker of several neurodegeneration diseases, we specifically focused on a subset of proteins genetically linked to neurodegeneration in humans. We mapped multiple ubiquitylation sites on neurodegeneration-associated proteins that consistently changed in mouse brain aging and following bortezomib treatment in 14 days iNeurons (Figure 3I, Table S4). This finding underlies that age-associated proteostasis impairment, e.g., decreased proteasome activity, can trigger ubiquitylation changes of proteins encoded by neurodegeneration-associated genes even in the absence of mutations.

## Discussion

In this study, we investigated the impact of aging on three commonly studied PTMs in the vertebrate brain. Our findings reveal that changes in protein ubiquitylation far exceed the signatures that can be detected from the same sample at the level of transcriptome, proteome, or other analyzed PTMs. Many of the ubiquitylation changes are independent of protein abundance and, if consistent, they show considerably more pronounced effect sizes. These differences cannot be attributed to technical artifacts related to peptide versus protein quantification, as other PTMs that undergo similar experimental (peptide enrichment) and analytical procedures do not exhibit such pronounced effects. Since ubiquitylation is a main regulator of protein degradation and homeostasis, one might expect that changes in this PTM would reflect in equal, if not greater, effects at the protein level. However, in line with previous observations (Koyuncu et al., 2021), our study shows this is not always the case. A possible explanation for the apparent paradox of changes of ubiquitylation that do not manifest in comparable protein level changes could reside in the fact that ubiquitylated proteoforms represent only a small percentage of the total protein pool (Prus et al., 2023), making their fluctuation in abundance less noticeable in comparison with the total protein pool. Alternatively, these sites might represent regulatory PTMs not involved in protein degradation. Nevertheless, we could observe a significant correlation between ubiquitylation and protein half-life changes (Kluever et al., 2022) during mouse brain aging (Figure 1I), suggesting a mechanistic link between altered protein turnover and accumulation of ubiquitylated proteoforms.

We describe a major remodeling of the protein ubiquitylation landscape, which is conserved in at least two vertebrate species (mouse and killifish). The remodeling of the protein ubiquitylation landscape with aging is in line with previous data from *C. elegans* (Koyuncu et al., 2021), and it demonstrates the conservation of this aspect of age-related proteostasis impairment in vertebrate species. Notably, an independent study reported that >90% of the proteins that become metastable in aging nematodes undergo changes of ubiquitylation (Sui et al., 2022), further underscoring a correlation between post-translational modification and loss of proteostasis.

In our study, many quantified ubiquitylation sites showed increased modification consistent with a global increase of protein ubiquitylation during brain aging (Gray et al., 2003; Matsui et al., 2019; Ohtsuka et al., 1995). Several chaperons and co-chaperons, e.g., DNAJB2, DNAJB6, and HSPA8, displayed pronounced increases of ubiquitylation both in mouse and killifish (Figure 1G-2C). HSPA8 has been reported to interact with tau fibrils (Baughman et al., 2018) and participate in selectively removing aggregated proteins in ALS (Crippa et al., 2010), while DNAJB1 and DNAJB6 are part of the DNAJ family, a group of co-chaperons that play a crucial role in regulating the activity of HSPA1 and assist in its proper functioning (Hennessy et al., 2005). These proteins might become hyper-ubiquitylated upon sequestration in protein aggregates (Walther et al., 2015).

Although most ubiquitylation sites show increased occupancy with aging, a subset decreased. Particularly, synaptic proteins were affected by a general decrease in ubiquitylation (Figure 1E), and previous reports showed that synaptic transmission is controlled by UPS (Tai et al., 2008). Interestingly, a substantial decrease in ubiquitylation was observed by (H. Chen et al., 2003) after the excitation of rat synaptosomes and was not reverted when blocking the proteasome, suggesting that deubiquitinases might regulate this process. Our data suggests that altered ubiquitylation of the synaptic compartment might play an essential role in how synaptic function and plasticity are altered in aging.

We experimentally assigned which changes in ubiquitylation levels are causally linked to decreased proteasome activity. We assessed this in human iNeurons treated with bortezomib, a proteasome inhibitor. Our analysis of the fraction of age-affected sites dependent on proteasome impairment is likely to be underestimated since we performed experiments in a single cell type (glutamatergic neurons). Therefore, our analysis might have missed ubiquitylation sites from proteins expressed by other cell types, or that did not align between mouse and human. Another potential limitation might arise from our iNeurons experiments comprising acute, relatively short treatments (24h) while aging cells experience a chronic reduction of proteasome activity. Nevertheless, a considerable part of the aging signature was recapitulated by the bortezomib treatment in iNeurons. Notably, the bafilomycin treatment showed a negative correlation, probably due to a compensatory increase of proteasome activity to counteract the lysosomal acidification impairment (Figure 3G).

Additional mechanisms might contribute to re-wiring the protein ubiquitylation landscape in the aging brain. Changes in the activity of deubiquitinases have been shown to influence protein ubiquitylation abundance and modulate organismal lifespan in nematodes (Koyuncu et al., 2021). Although we did not measure the activity of these enzymes directly, we found a subset of proteins involved in the ubiquitin cycle to be significantly changing in abundance during brain aging (Figure S4D). It is tempting to speculate that the altered activity of these enzymes might contribute to age-related protein ubiquitylation changes in vertebrates.

Finally, we identified ubiquitylation sites in the brain for several proteins genetically linked to human neurodegeneration. Importantly, we found multiple of these sites to be affected by aging (Figure 3I and Table S4). Although it remains unclear whether these PTMs are mechanistically linked to the pathogenic properties of the mutated forms of these proteins or whether the same modifications occur in diseased human tissues, our findings suggest that ubiquitylated proteoforms are generally more susceptible to age-related changes and may represent more robust biomarkers of brain aging and, potentially, neurodegeneration.

## Supplementary figures

**Figure S1.**
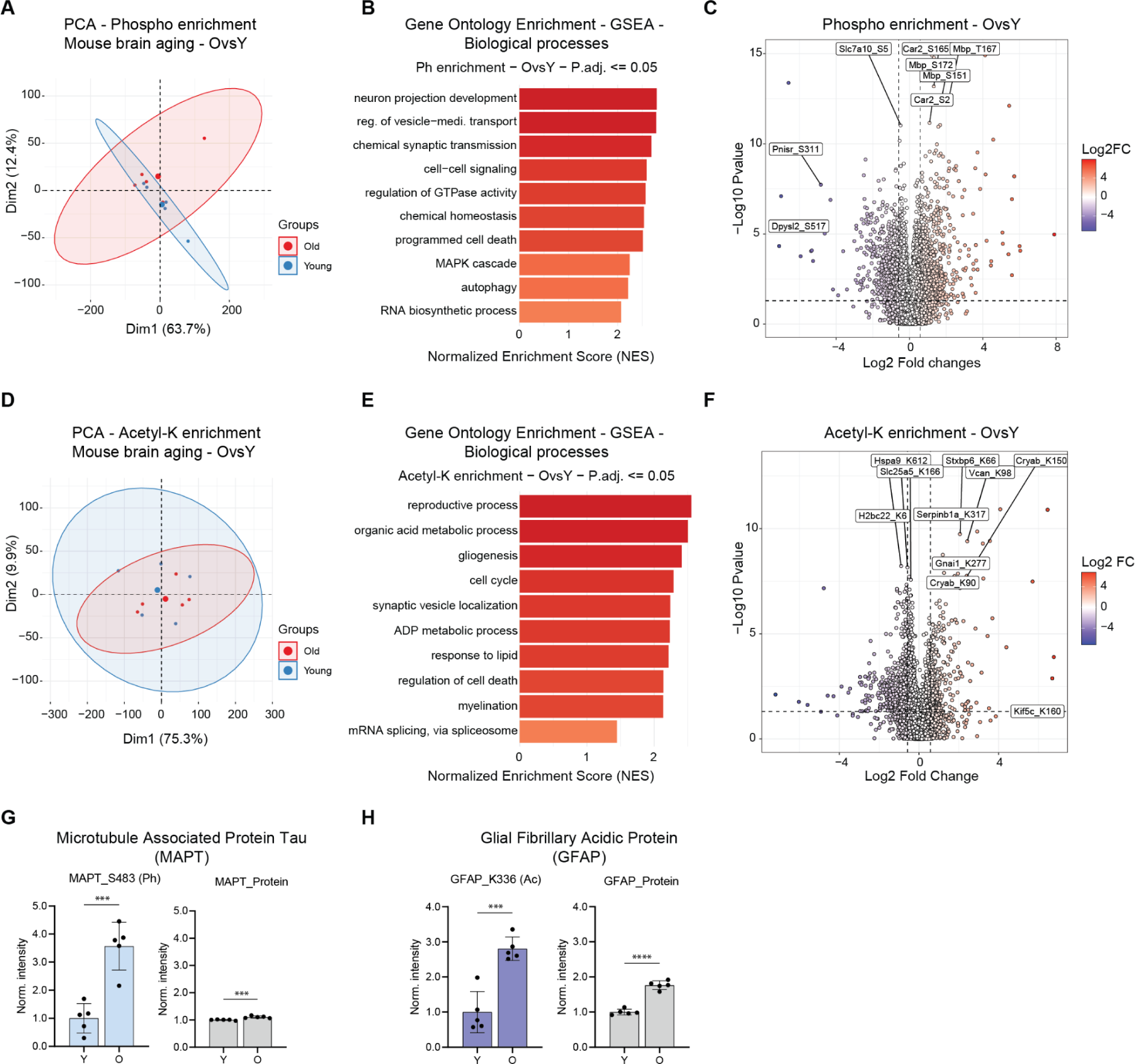
Phosphorylation and Acetylation Characterization in the Mouse Aging Brain. A) PCA plot of phosphorylation changes in mouse brain aging. Ellipses represent 95% confidence intervals. The percentage of variance explained by each principal component is indicated. B) GO enrichment analysis for phospho-enrichment using GSEA Biological processes (Adj.PValue<0.05). C) Volcano plot for phospho-enrichment in mouse brain aging. D) PCA plot of acetylation changes in mouse brain aging. Ellipses represent 95% confidence intervals. The percentage of variance explained by each principal component is indicated. E) GO enrichment analysis for acetyl-enrichment using GSEA Biological processes (Adj.PValue<0.05) F) Volcano plot for acetyl-enrichment in mouse brain aging. G) MAPT phosphorylation during aging, in light blue, the phosphorylation level of the serine 483, in gray change in total MAPT protein level (N=5, unpaired *t*-test). H) GFAP acetylation during aging, in dark blue, the acetylation level of the lysine 336; in gray, change in total MAPT protein level (N=5, unpaired *t*-test). *P ≤ 0.05; **P ≤ 0.01, ***P ≤ 0.001, ****P ≤ 0.0001.

**Figure S2.**
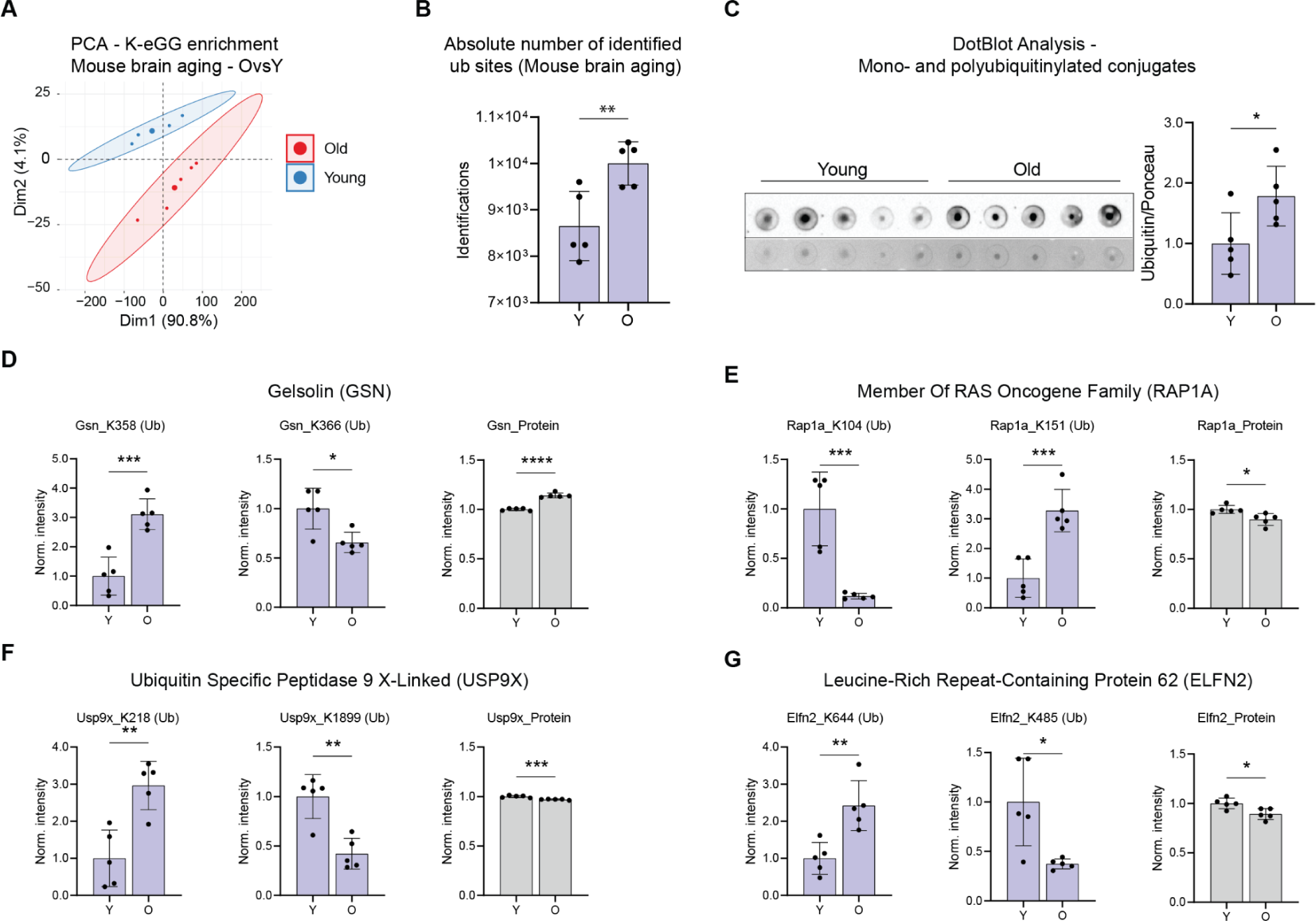
Changes in ubiquitylation in mouse brain aging. A) PCA plot of ubiquitylation changes in mouse brain aging. Ellipses represent 95% confidence intervals. The percentage of variance explained by each principal component is indicated. B) Barplots showing the absolute numbers of identified ubiquitylated sites in the different age groups (N=5, unpaired *t*-test). C) Dot Blot analysis of total ubiquitin conjugates and free ubiquitin in young and old mouse brain lysates on the right barplot with the signal quantification (N=5, unpaired *t*-test). D) Examples of proteins showing both an increase and decrease level of ubiquitylation (N=5, unpaired *t*-test). *P ≤ 0.05; **P ≤ 0.01, ***P ≤ 0.001, ****P ≤ 0.0001.

**Figure S3.**
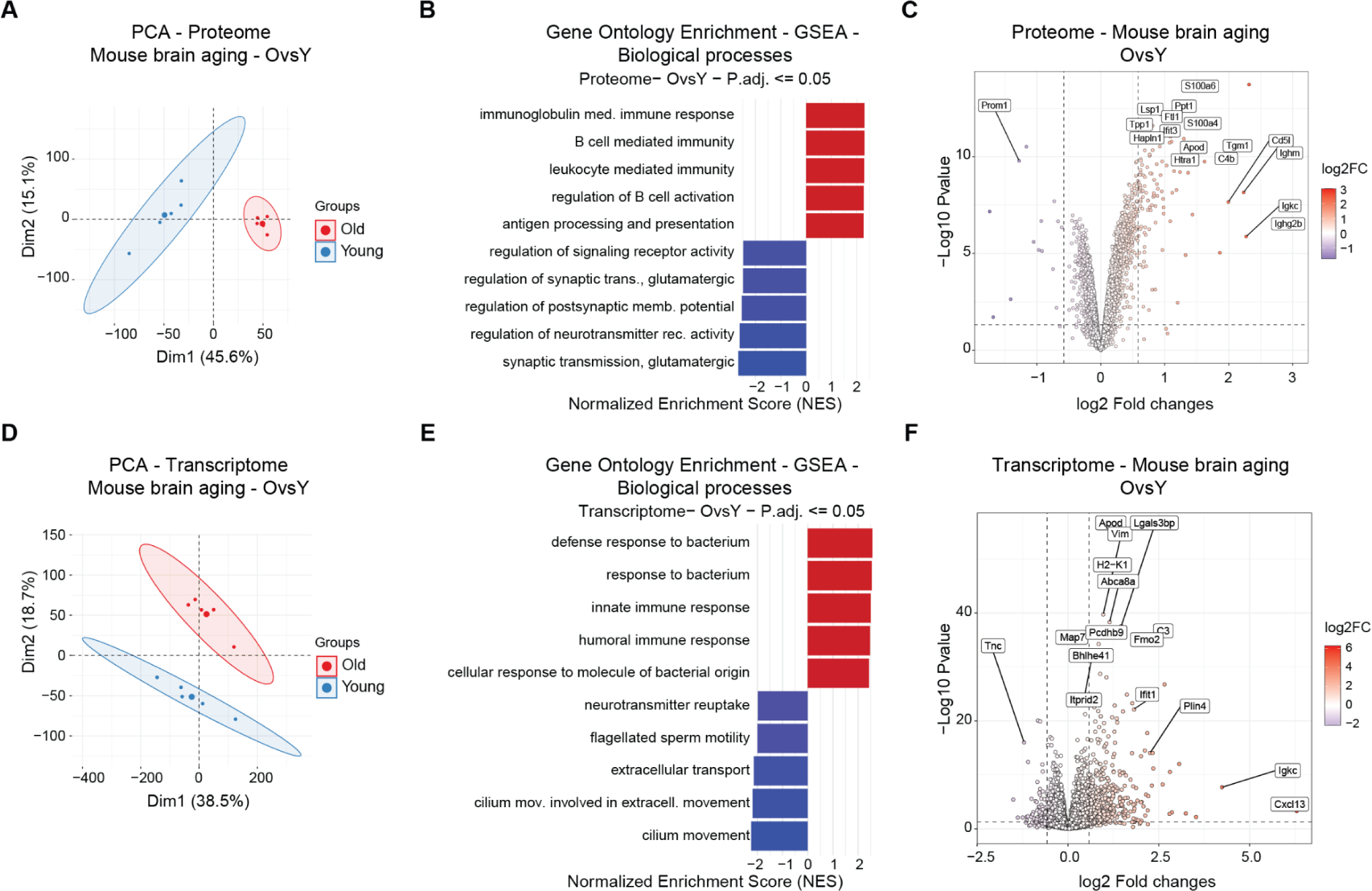
Changes in whole proteome and transcriptome in mouse brain aging. A) PCA plot of whole proteome changes in mouse brain aging. Ellipses represent 95% confidence intervals. The percentage of variance explained by each principal component is indicated. B) GO enrichment analysis for proteome using GSEA Biological processes (Adj.PValue<0.05). C) Volcano plot for proteome in mouse brain aging (N=5). D) PCA plot of transcriptome changes in mouse brain aging. Ellipses represent 95% confidence intervals. The percentage of variance explained by each principal component is indicated. E) GO enrichment analysis for transcriptome using GSEA Biological processes (Adj.PValue<0.05). F) Volcano plot for the transcriptome in mouse brain aging (N=5). *P ≤ 0.05; **P ≤ 0.01, ***P ≤ 0.001, ****P ≤ 0.0001.

**Figure S4.**
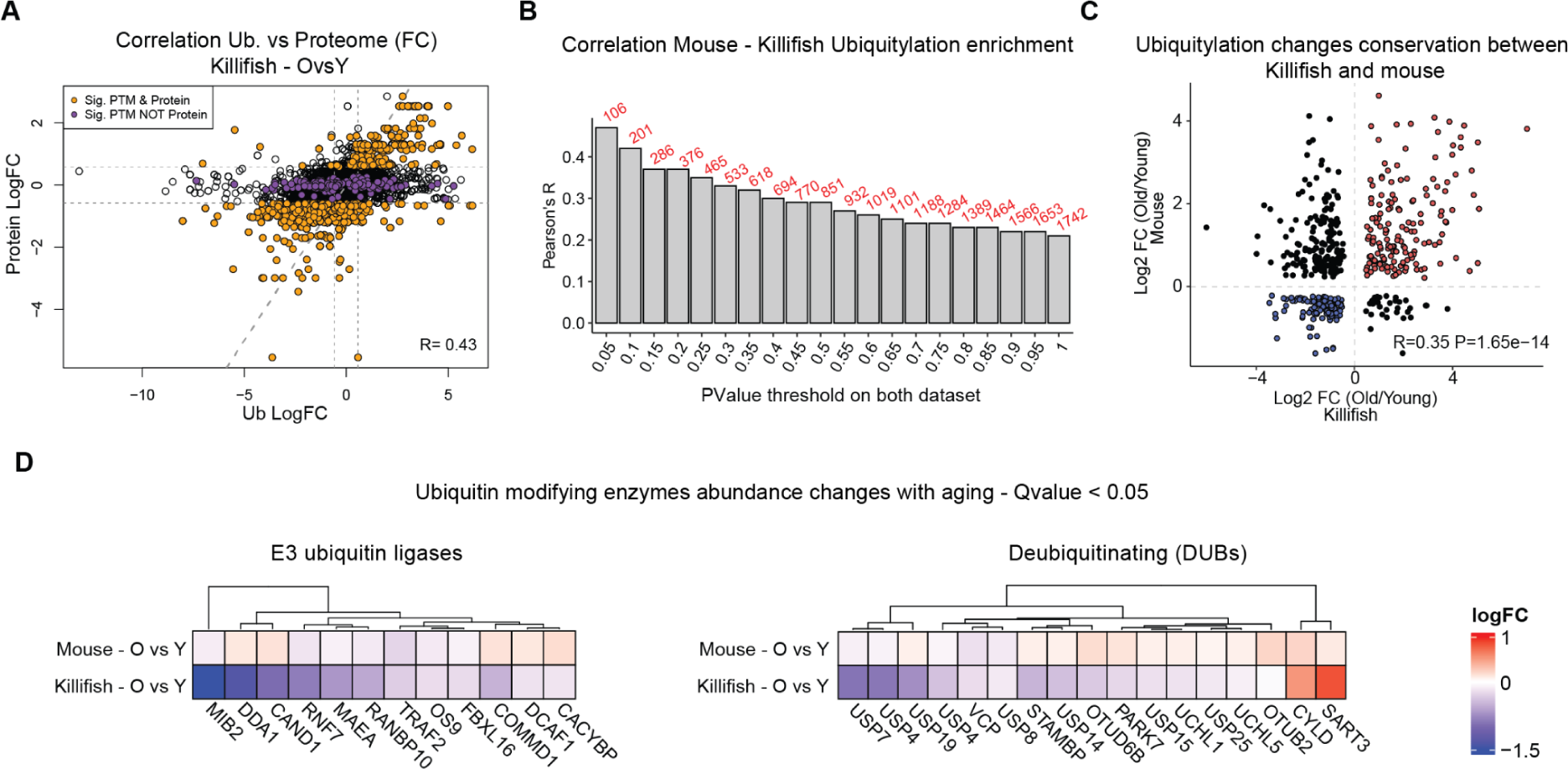
Ubiquitylation changes conservation during vertebrate brain aging. A) Scatterplot describing the correlation between proteome and ubiquitylome, in orange are highlighted proteins which are affected significantly in both layers (Ub QValue<0.05; proteome Adj.PValue<0.05 and FC>0.58 & <-0.58), in purple proteins which are significantly changing only for ubiquitylation (Ub QValue<0.05; proteome Adj.PValue>0.05). B) Barplots showing PearsońR value on the y-axis for mouse and killifish ubiquitylomes correlation. On the x-axis, different PValue cutoff thresholds are applied. In red numbers of modified sites that pass the respective thresholds. C) Scatterplot for ubiquitylome comparison between mouse and killifish. In red are highlighted the aligned modified sites that increase ubiquitylation in both species, and in blue are highlighted the ones that decrease (PValue<0.25 on both datasets). D) Heatmap showing protein abundance change of E3 ubiquitin ligases (upper panel) and deubiquitinating enzymes (bottom panel) in mouse and killifish during aging (QValue cutoff for both datasets <0.05). *P ≤ 0.05; **P ≤ 0.01, ***P ≤ 0.001, ****P ≤ 0.0001.

**Figure S5.**
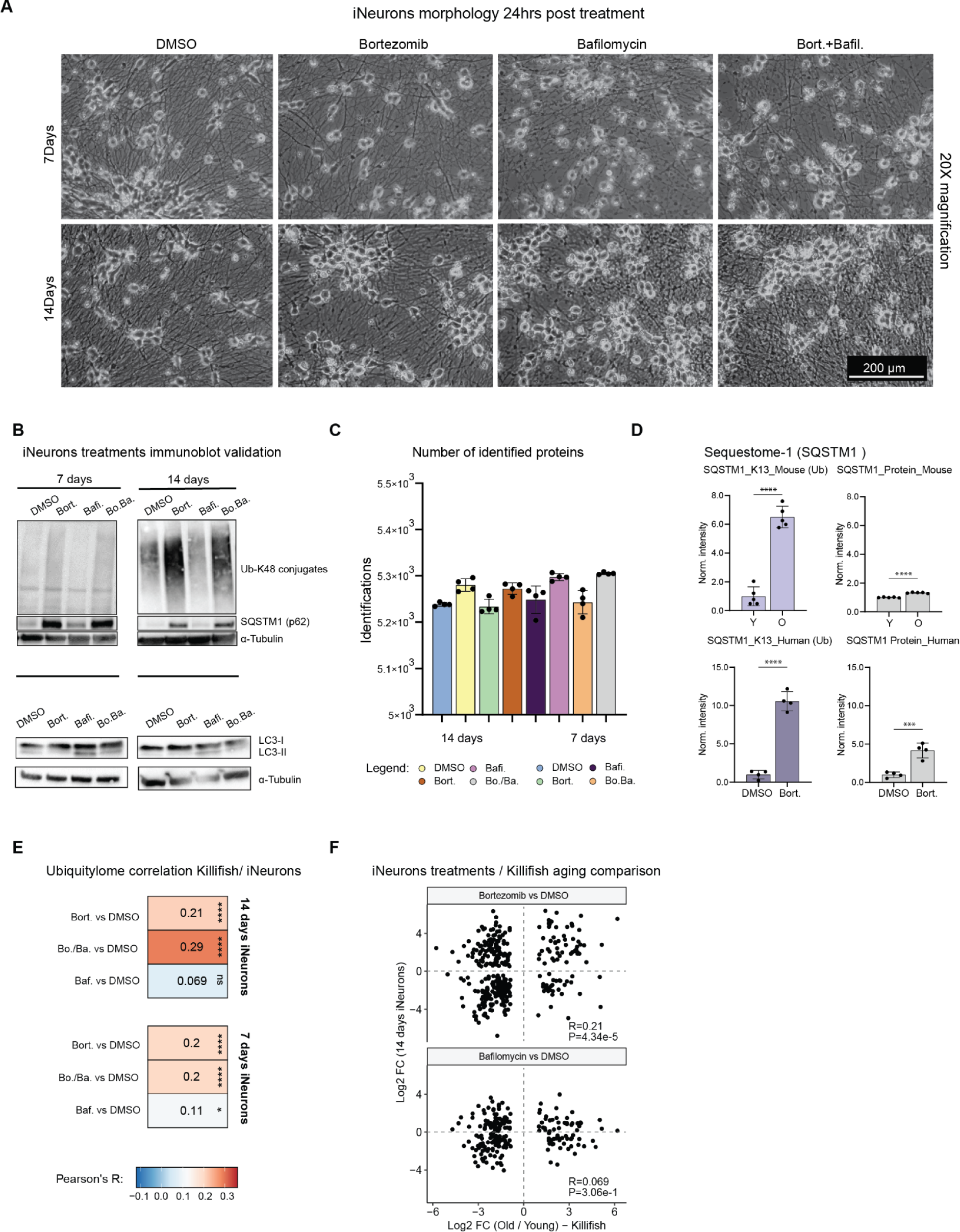
Proteasome inhibition in iNeurons and comparison with the aging datasets. A) Representative pictures of 7 days and 14 days iNeurons cell morphology 24 hours post-treatment (20X magnification, scale 200 µm). B) Immunoblot validation of bortezomib, bafilomycin, and combination of both in iNeurons. For the bortezomib treatment is visible an increase in K-48 ubiquitylated conjugates together with an increase in SQSTM1 (p62) levels; for the bafilomycin treatment, an increase in LC3-II/LC3-I is visible. C) Barplots showing the absolute number of identified proteins in the different iNeurons sample groups. D) SQSTM1 (p62) ubiquitylation, during aging (light purple) and upon proteasome inhibition in 14 days iNeurons (dark purple), in gray change in total SQSTM1 protein level (N=5 for mouse and N=4 for iNeurons, unpaired *t*-test). E) Correlation between iNeurons treatment and killifish aging datasets. F) Scatterplot for ubiquitylome comparison between mouse aging and 14 days iNeurons treated with bortezomib (upper panel) or bafilomycin (lower panel) compared to DMSO. *P ≤ 0.05; **P ≤ 0.01, ***P ≤ 0.001, ****P ≤ 0.0001.

**Figure S6.**
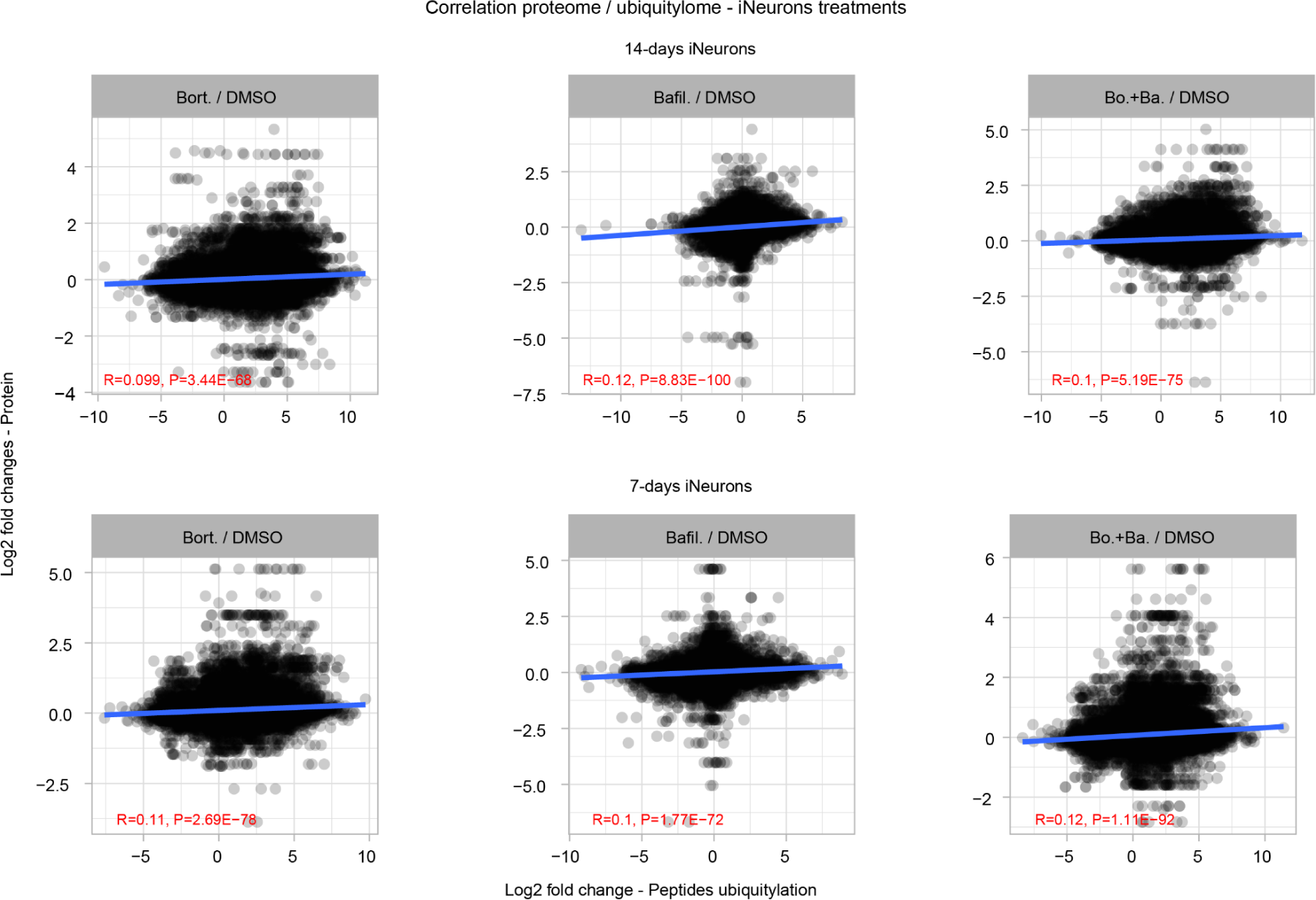
iNeurons proteome / ubiquitylomes correlation in different treatments. Relationship between iNeurons treatments abundance changes of modified peptides vs. corresponding protein. The red text indicates the test results for the association between paired samples using Pearson’s product-moment correlation coefficients.

**Figure S7.**
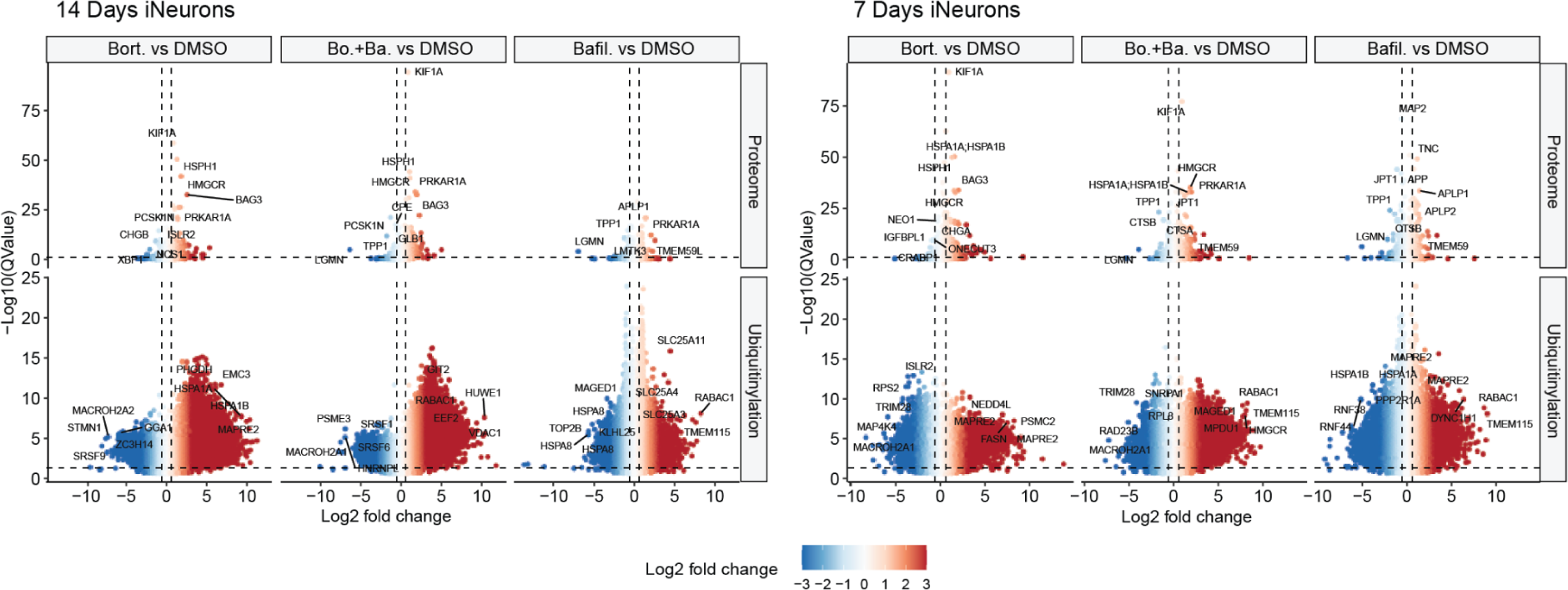
iNeurons ubiquitylomes changes exceed proteome changes. Volcano plots depicting the changes in the whole proteome and ubiquitylome of iNeurons upon treatment with bortezomib, bafilomycin, and a combination of both drugs. The upper panel represents proteome data, while the lower lane represents ubiquitylome data (N=4, horizontal dashed line shows QValue=0.05 while vertical lines show fold changes = –0.58, 0.58).

## Acknowledgments

The authors gratefully acknowledge support from the FLI Core Facilities Proteomics, and the Fish and Mouse Facilities. A.O. is supported by the German Research Council (Deutsche Forschungsgemeinschaft, DFG) via the Research Training Group ProMoAge (GRK 2155), the Else Kröner Fresenius Stiftung (award number: 2019_A79), the Fritz-Thyssen Foundation (award number: 10.20.1.022MN), the Chan Zuckerberg Initiative Neurodegeneration Challenge Network (award numbers: 2020-221617, 2021-230967 and 2022-250618), and the NCL Stiftung. The FLI is a member of the Leibniz Association and is financially supported by the Federal Government of Germany and the State of Thuringia.

## Author contributions

Conceptualization: AM, DDF, AO

Data curation: AM, DDF

Investigation: AM, DP, AKS

Methodology: AM

Project administration: AO

Data analysis: AM, DDF

Supervision: AO

Visualization: AM, DDF

Writing – original draft: AM, DDF, AO

Writing – review & editing: DP

## Material and methods

### Mice

All wild-type mice were C57BL/6J obtained from Janvier Labs (Le Genest-Saint-Isle, France) or internal breeding at FLI. All animals were kept in a specific pathogen-free animal facility with a 12 h light/dark cycle at a temperature of 20 °C ± 2 and humidity of 55% ± 15. Young mice were aged three months, and old mice were aged 33 months. During the experiment, Mice had unlimited access to food (ssniff, Soest, Germany). For the proteomic and transcriptomic analysis, only male mice were used. Mice were euthanized with CO_2_, and organs were isolated, washed in PBS, weighted, and immediately snap-frozen in liquid nitrogen before storage at −80 °C. All experiments were carried out according to the guidelines from Directive 2010/63/EU of the European Parliament on the protection of animals used for scientific purposes. The protocols of animal maintenance and euthanasia were approved by the local authorities for animal welfare in the State of Thuringia, Germany.

### hiPSC Cell Line maintenance and differentiation

The protocol used for the maintenance of the WTC11 hiPSC cell line is described in detail by (Wang et al., 2017). In brief, the Matrigel (Corning, 356231) coating solution was prepared by diluting 100X concentrated Matrigel Matrix in DMEM/F12 (Gibco, 11320033) medium and coating the tissue culture surface with it. Thawed cells were suspended in Essential 8 (E8) culture medium (Gibco, A1517001) supplemented with Y-27632 ROCK inhibitor (Abcam, ab120129) and transferred to the Matrigel-coated plate. The culture was maintained by daily E8 medium change without ROCK inhibitor. Cells were split using Accutase solution (Sigma, A6964) and resuspended in prewarmed E8 culture medium supplemented with ROCK inhibitor before transfer to the Matrigel-coated plate. For long-term storage, cells were frozen in cryopreservation medium (10% DMSO, 20% FBS in E8 medium) and transferred to liquid nitrogen. WTC11 hiPSC Cell Line was a kind gift from Ward Lab.

### iPSCs differentiation into iNeurons and drug treatments

Cells were collected using Accutase solution (Sigma, A6964) when they reached 70% – 80% confluency. After centrifugation, the cell pellet was resuspended and transferred to a Matrigel-coated plate. The induction medium (IM) (DMEM/F12 (Gibco, 11320033) supplemented with N2 supplement (Gibco, 17502048), MEM NEAA (Gibco, 11140050), L-glutamine (Gibco, 25030081) and 2 µg/ml doxycycline (Sigma, D9891)) supplemented with 10 µM Y-27632 ROCK inhibitor (Abcam, ab120129) was then added to the cells, which were incubated at 37 °C overnight. The IM without ROCK inhibitor was exchanged on the second day, and on the third day, cells were collected using Accutase solution and either frozen or re-plated for neuronal maturation. To culture iNeurons, PLO-coating was performed (1 mg/ml Poly-L-ornithine hydrobromide (Sigma Aldrich, 27278-49-0) in the buffer containing 100 mM boric acid, 25 mM sodium tetraborate, 75 mM sodium chloride and 1 M sodium hydroxide), and freshly dissociated 3-day differentiated iNeurons were resuspended in cortical neuron culture medium (CM) (Neurobasal medium (Gibco, 21103049) supplemented with B-27 supplement (Gibco, 17504044), 10 ng/ml BDNF (PeproTech, 450-02), 10 ng/ml NT-3 (PeproTech, 450-03) and 1 µg/ml laminin (Sigma-Aldrich, 114956-81-9) supplemented with 2 µg/ml doxycycline and 10 µM Y-27632 ROCK inhibitor. Half-medium changes with CM were performed biweekly to maintain the culture.

The proteasome and lysosome acidification inhibitions were performed in either 7-days or 14-days iNeurons. Proteasome inhibition was achieved by treating cells with bortezomib (Sigma, 5043140001), and lysosome acidification was inhibited using Bafilomycin A1 from *S. griseus* (Sigma, B1793). Vehicle solution (DMSO) and 20 nM drug solutions were prepared in one-half culture volume of fresh CM. Half of the culture medium was removed from the cell culture plate directly before treatment and substituted with an equal amount of corresponding, fresh-prepared drug solution, resulting in a 10 nM final concentration of the drug. Plates were then returned to the incubator and incubated for 24 hours.

### Sample preparation for total proteome and analysis of PTMs

For iNeurons samples, cell pellets were used directly for the lysis step. For snap-frozen brains, samples were thawed and transferred into Precellys® lysing kit tubes (Keramik-kit 1.4/2.8 mm, 2 ml (CKM)) containing PBS supplemented with cOmplete™, Mini, EDTA-free Protease Inhibitor (Roche,11836170001) and with PhosSTOP™ Phosphatase Inhibitor (Roche, 4906837001). The volume of PBS added was calculated based on an estimated protein content (5% of fresh tissue weight) to reach a 10 µg/µl concentration. Tissues were homogenized twice at 6000 rpm for 30 s using Precellys® 24 Dual (Bertin Instruments, Montigny-le-Bretonneux, France), and the homogenates were transferred to new 2 ml Eppendorf tubes.

Samples corresponding to ∼1.5 mg of protein were used as starting material for each biological replicate. Volumes were adjusted using PBS and samples were lysed by the addition of 4× lysis buffer (8% SDS, 100 mM HEPES, pH8). Samples were sonicated twice in a Bioruptor Plus for 10 cycles with 1 min ON and 30 s OFF with high intensity at 20 °C. The lysates were centrifuged at 18,407 x*g* for 1 min and transferred to new 1.5 ml Eppendorf tubes. Subsequently, samples were reduced using 20 mM DTT (Carl Roth, 6908) for 15 min at 45 °C and alkylated using freshly made 200 mM iodoacetamide (IAA) (Sigma-Aldrich, I1149) for 30 min at room temperature in the dark. An aliquot of each lysate was used for estimating the precise protein quantity using Qubit protein assay, and volumes were adjusted accordingly to have a total protein amount of 1.25 mg (Thermo Scientific, Q33211). Subsequently, proteins were precipitated using cold acetone, as described in (Buczak et al., 2020), and resuspended in 500 µl of digestion buffer (3 M urea, 100 mM HEPES pH 8.0). Proteins were digested using LysC 1:100 enzyme:proteins ratio for 4 hours (Wako sequencing grade, 125-05061) and trypsin 1:100 enzyme:proteins ratio for 16 hours (Promega sequencing grade, V5111). The digested proteins were then acidified with 10% (v/v) trifluoroacetic acid. Aliquots corresponding to 20, 200, and 1000 µg of peptides were taken for proteome, phosphopeptides, and ubiquitylated / acetylated peptides enrichment, respectively, and desalted using Waters Oasis® HLB µElution Plate 30 µm (2, 10, and 30 mg, depending on the amount of starting material) following manufacturer instructions. The eluates were dried down using a vacuum concentrator and reconstituted in MS buffer A (5% (v/v) acetonitrile, 0.1% (v/v) formic acid). For PTM enrichment, peptides were further processed as described below. For Data Independent Acquisition (DIA) based analysis of total proteome, samples were transferred to MS vials, diluted to a concentration of 1 µg/µL, and spiked with iRT kit peptides (Biognosys, Ki-3002-2) before analysis by LC-MS/MS.

### Sequential enrichment of ubiquitylated and acetylated peptides

Ubiquitylated and acetylated peptides were sequentially enriched starting from ∼1000 µg of dried peptides per replicate. For the enrichment of ubiquitylated peptides, the PTMScan® HS Ubiquitin/SUMO Remnant Motif (K-ε-GG) kit (Cell Signaling Technology, 59322) was used following manufacturer instructions. The K-ε-GG modified enriched fraction was desalted and concentrated as described above, dissolved in MS buffer A, and spiked with iRT kit peptides before LC-MS/MS analysis.

The flowthrough fractions from the K-ε –GG enrichment were acidified with 10% (v/v) trifluoroacetic acid and desalted using Oasis® HLB µElution Plate 30 µm (30 mg) following manufacturer instructions. Acetylated peptides were enriched as described by (Di Sanzo et al., 2021). Briefly, dried peptides were dissolved in 1000 µl of IP buffer (50 mM MOPS pH 7.3, 10 mM KPO_4_ pH 7.5, 50 mM NaCl, 2.5 mM Octyl β-D-glucopyranoside) to reach a peptide concentration of 1 µg/µL, followed by sonication in a Bioruptor Plus (5 cycles with 1 min ON and 30 s OFF with high intensity at 20 °C). Agarose beads coupled to an antibody against acetyl-lysine (ImmuneChem Pharmaceuticals Inc., ICP0388-5MG) were washed three times with washing buffer (20 mM MOPS pH 7.4, 10 mM KPO4 pH 7.5, 50 mM NaCl) before incubation with each peptide sample for 1.5 h on a rotating well at 750 rpm (STARLAB Tube roller Mixer RM Multi-1). Samples were transferred into Clearspin filter microtubes (0.22 µm) (Dominique Dutscher SAS, Brumath, 007857ACL) and centrifuged at 4 °C for 1 min at 2000 x*g*. Beads were washed first with IP buffer (three times), then with washing buffer (three times), and finally with 5 mM ammonium bicarbonate (three times). Thereupon, the enriched peptides were eluted first in basic condition using 50 mM aqueous NH3, then using 0.1% (v/v) trifluoroacetic acid in 10% (v/v) 2-propanol and finally with 0.1% (v/v) trifluoroacetic acid. Elutions were dried down and reconstituted in MS buffer A (5% (v/v) acetonitrile, 0.1% (v/v) formic acid), acidified with 10% (v/v) trifluoroacetic acid, and then desalted with Oasis® HLB µElution Plate 30 µm. Desalted peptides were finally dissolved in MS buffer A, spiked with iRT kit peptides, and analyzed by LC-MS/MS.

### Enrichment of phosphorylated peptides

Desalted peptides corresponding to 200 µg, as described in ‘‘Sample preparation for total proteome and analysis of PTMs’ were used. The last desalting step was performed using 50 μl of 80% ACN and 0.1% TFA buffer solution. Before phosphopeptide enrichment, samples were filled up to 210 µl using 80% ACN and 0.1% TFA buffer solution. Phosphorylated peptides were enriched using Fe(III)-NTA cartridges (Agilent Technologies, G5496-60085) in an automated fashion using the standard protocol from the AssayMAP Bravo Platform (Agilent Technologies). In short, Fe(III)-NTA cartridges were first primed with 100 µl of priming buffer (100% ACN, 0.1% TFA) and equilibrated with 50 μL of buffer solution (80% ACN, 0.1% TFA). After loading the samples into the cartridge, the cartridges were washed with an OASIS elution buffer, while the syringes were washed with a priming buffer (100% ACN, 0.1% TFA). The phosphopeptides were eluted directly with 25 μL of 1% ammonia into 25 μL of 10% FA. Samples were dried with a speed vacuum centrifuge and stored at −20 °C until LC-MS/MS analysis.

### Data-independent acquisition

Peptides were separated in trap/elute mode using the nanoAcquity MClass Ultra-High Performance Liquid Chromatography system (Waters, Waters Corporation, Milford, MA, USA) equipped with trapping (nanoAcquity Symmetry C18, 5 μm, 180 μm × 20 mm) and an analytical column (nanoAcquity BEH C18, 1.7 μm, 75 μm × 250 mm). Solvent A was water and 0.1% formic acid, and solvent B was acetonitrile and 0.1% formic acid. 1 μl of the samples (∼1 μg on column) were loaded with a constant flow of solvent A at 5 μl/min onto the trapping column. Trapping time was 6 min. Peptides were eluted via the analytical column with a constant flow of 0.3 μl/min. During the elution, the percentage of solvent B increased nonlinearly from 0–40% in 120 min. The total run time was 145 min, including equilibration and conditioning. The LC was coupled to an Orbitrap Exploris 480 (Thermo Fisher Scientific, Bremen, Germany) using the Proxeon nanospray source. The peptides were introduced into the mass spectrometer via a Pico-Tip Emitter 360-μm outer diameter × 20-μm inner diameter, 10-μm tip (New Objective) heated at 300 °C, and a spray voltage of 2.2 kV was applied. The capillary temperature was set at 300°C. The radio frequency ion funnel was set to 30%. For DIA data acquisition, full scan mass spectrometry (MS) spectra with a mass range 350–1650 m/z were acquired in profile mode in the Orbitrap with a resolution of 120,000 FWHM. The default charge state was set to 3+. The filling time was set at a maximum of 60 ms with a limitation of 3 × 10^6^ ions. DIA scans were acquired with 40 mass window segments of differing widths across the MS1 mass range. Higher collisional dissociation fragmentation (stepped normalized collision energy; 25, 27.5, and 30%) was applied, and MS/MS spectra were acquired with a resolution of 30,000 FWHM with a fixed first mass of 200 m/z after accumulation of 3 × 10^6^ ions or after filling time of 35 ms (whichever occurred first). Data were acquired in profile mode. Xcalibur 4.3 (Thermo) and Tune version 2.0 were used to acquire and process the raw data.

### TMT labeling

TMT was used for mouse proteome data. The solution containing the resuspended peptides were brought to a pH 8.5 and a final concentration of 100 mM HEPES (Sigma H3375) prior to labeling. 20 mg of peptides were used for each label reaction. TMT-10plex reagents (Thermo Fisher #90111) were reconstituted in 41 mL of acetonitrile (Biosolve #0001204102BS). TMT labeling was performed in two steps by addition of 2x of the TMT reagent per mg of peptide (e.g., 40 mg of TMT reagent for 20 mg of peptides). TMT reagents were added to samples at room temperature, followed by incubation in a thermomixer (Eppendorf) under constant shaking at 600 rpm for 30 min. After incubation, a second portion of TMT reagent was added and followed by incubation for another 30 min. After checking the labeling efficiency by MS, equal amounts of samples were pooled (200 mg total), desalted using two wells of a Waters Oasis HLB mElution Plate 30 mm (Waters #186001828BA) and subjected to high pH fractionation prior to MS analysis. The labeling of the samples was performed as follows:

|  |  |  |  |  |  |  |  |  |  |
| --- | --- | --- | --- | --- | --- | --- | --- | --- | --- |
| Young1 | Young2 | Young3 | Young4 | Young5 | Old1 | Old2 | Old3 | Old4 | Old5 |
| 126 | 127_N | 127_C | 128_N | 128_C | 129_N | 129_C | 130_N | 130_C | 131 |

### High pH peptide fractionation

Offline high pH reverse phase fractionation was performed using an Agilent 1260 Infinity HPLC System equipped with a binary pump, degasser, variable wavelength UV detector (set to 220 and 254 nm), peltier-cooled autosampler (set at 10C) and a fraction collector. The column used was a Waters XBridge C18 column (3.5 mm, 100 3 1.0 mm, Waters) with a Gemini C18, 4 3 2.0 mm SecurityGuard (Phenomenex) cartridge as a guard column. The solvent system consisted of 20 mM ammonium formate (20 mM formic acid (Biosolve #00069141A8BS), 20 mM (Fluka #9857) pH 10.0) as mobile phase (A) and 100% acetonitrile (Biosolve #0001204102BS) as mobile phase (B). The separation was performed at a mobile phase flow rate of 0.1 mL/min using a non-linear gradient from 95% A to 40% B for 91 min. Forty-eight fractions were collected along with the LC separation that were subsequently pooled into 23 or 24 fractions. Pooled fractions were dried in a speed vacuum centrifuge and then stored at –80 °C until MS analysis.

### Data acquisition for TMT labeled samples

For TMT experiments, fractions were resuspended in 20 mL reconstitution buffer (5% (v/v) acetonitrile (Biosolve #0001204102BS), 0.1% (v/v) TFA in water) and 5 mL were injected into the mass spectrometer. Peptides were separated using the nanoAcquity UPLC system (Waters) fitted with a trapping (nanoAcquity Symmetry C18, 5 mm, 180 mm x 20 mm) and an analytical column (nanoAcquity BEH C18, 2.5 mm, 75 mm x 250 mm). The analytical column outlet was coupled directly to an Orbitrap Fusion Lumos (Thermo Fisher Scientific) using the Proxeon nanospray source. Solvent A was water with 0.1% (v/v) formic acid and solvent B was acetonitrile, 0.1% (v/v) formic acid. The samples were loaded with a constant flow of solvent A at 5 ml/min, onto the trapping column. Trapping time was 6 min. Peptides were eluted via the analytical column at a constant flow rate of 0.3 ml/ min, at 40C. During the elution step, the percentage of solvent B increased in a linear fashion from 5% to 7% in the first 10 min, then from 7% B to 30% B in the following 105 min and to 45% B by 130 min. The peptides were introduced into the mass spectrometer via a Pico-Tip Emitter 360 mm OD x 20 mm ID; 10 mm tip (New Objective) and a spray voltage of 2.2kV was applied. The capillary temperature was set at 300C. Full scan MS spectra with a mass range of 375-1500 m/z were acquired in profile mode in the Orbitrap with a resolution of 60000 FWHM using the quad isolation. The RF on the ion funnel was set to 40%. The filling time was set to a maximum of 100 ms with an AGC target of 4 3 105 ions and 1 microscan. The peptide monoisotopic precursor selection was enabled along with relaxed restrictions if too few precursors were found. The most intense ions (instrument operated for a 3 s cycle time) from the full scan MS were selected for MS2, using quadrupole isolation and a window of 1 Da. HCD was performed with a collision energy of 35%. A maximum fill time of 50 ms for each precursor ion was set. MS2 data were acquired with a fixed first mass of 120 m/z and acquired in the ion trap in Rapid scan mode. The dynamic exclusion list was set with a maximum retention period of 60 s and a relative mass window of 10 ppm. For the MS3, the precursor selection window was set to the range 400-2000 m/z, with an exclusion width of 18 m/z (high) and 5 m/z (low). The most intense fragments from the MS2 experiment were co-isolated (using Synchronus Precursor Selection = 8) and fragmented using HCD (65%). MS3 spectra were acquired in the Orbitrap over the mass range of 100-1000 m/z and the resolution set to 30000 FWHM. The maximum injection time was set to 105 ms and the instrument was set not to inject ions for all available parallelizable times. For data acquisition and processing of raw data the Xcalibur v4.0 and Tune v2.1 were used.

### Immunoblot

iNeurons or mouse brains were lysed as described in “Sample preparation for total proteome and analysis of PTMs.” Protein concentration was estimated by Qubit assay (Invitrogen, Q33211), and 30 µg of proteins were used. 4× loading buffer (1.5 M Tris pH 6.8, 20% (w/v) SDS, 85% (v/v) glycerin, 5% (v/v) β-mercaptoethanol) was added to each sample and then incubated at 95 °C for 5 minutes. Proteins were separated on 4–20% Mini-Protean® TGX™ Gels (BioRad, 4561096) by sodium dodecyl sulfate-polyacrylamide gel electrophoresis (SDS-PAGE) using a Mini-Protean® Tetra Cell system (BioRad, Neuberg, Germany, 1658005EDU).

Proteins were transferred to a nitrocellulose membrane (Carl Roth, 200H.1) using a Trans-Blot® Turbo™ Transfer Starter System (BioRad, 1704150). Membranes were stained with Ponceau S (Sigma, P7170-1L) for 5 min on a shaker (Heidolph Duomax 1030), washed with Milli-Q water, imaged on a Molecular Imager ChemiDocTM XRS + Imaging system (BioRad) and destained by 2 washes with PBS and 2 washes in TBST (Tris-buffered saline (TBS, 25 mM Tris, 75 mM NaCl), with 0.5% (v/v) Tween-20) for 5 min. After incubation for 5 min in EveryBlot blocking buffer (Biorad, 12010020), membranes were incubated overnight with primary antibodies against total ubiquitin-conjugates (Enzo Life Sciences, BML-PW8810), SQSTM1 (Abcam, ab91526), ubiquitin lys48-Specific (MilliporeSigma, 05-1307), LC3 (Cell Signaling Technology, 2775) or α-tubulin (Sigma, T9026) diluted (1:1000) in enzyme dilution buffer (0.2% (w/v) BSA, 0.1% (v/v) Tween20 in PBS) at 4 °C on a tube roller (BioCote® Stuart® SRT6). Membranes were washed 3 times with TBST for 10 min at room temperature and incubated with horseradish peroxidase coupled secondary antibodies (Dako, P0448/P0447) at room temperature for 1 h (1:2000 in 0.3% (w/v) BSA in TBST). After 3 more washes for 10 min in TBST, chemiluminescent signals were detected using ECL (enhanced chemiluminescence) Pierce detection kit (Thermo Fisher Scientific, Waltham, MA, USA, #32109). Signals were acquired on the Molecular Imager ChemiDocTM XRS + Imaging system and analyzed using the Image Lab 6.1 software (Biorad). Membranes were stripped using stripping buffer (1% (w/v) SDS, 0.2 M glycine, pH 2.5), washed 3 times with TBST, blocked, and incubated with the second primary antibody, if necessary.

### RNA isolation for RNA-Seq analysis

Individual brains from the mice were collected and snap-frozen in liquid nitrogen. On the day of the experiment, samples were thawed on ice, 1 mL of ice-cold Qiazol (Qiagen, 79306) reagent was added and transferred into Precellys® lysing kit tubes (Keramik-kit 1.4/2.8 mm, 2 ml (CKM)) Tissues were homogenized twice at 6000 rpm for 30 s using Precellys® 24 Dual (Bertin Instruments, Montigny-le-Bretonneux, France), transferred to new 2 ml Eppendorf tubes and then further processed using 1 mL syringes with 26G needles. The RNA extraction was carried out according to the manufacturer’s instructions. RNA integrity number was checked at the end of the procedure. Samples were further processed by GENEWIZ from Azenta Life Sciences from the library preparation to the bioinformatic analysis. RNA-seq was performed using Illumina NovaSeq 2×150 bp sequencing, 10M read pairs, and PolyA selection with ERCC spike-in.

## Data analysis

### Data processing for mass spectrometry DIA data

Spectral libraries were created for the PTMs by searching DIA and/or DDA runs using Spectronaut Pulsar (18.2.33, Biognosys, Zurich, Switzerland), while iNeurons proteome library was generated with Spectronaut Pulsar vr. 16.2.22, as referred to in Table 1. The data were searched against species-specific protein databases with an appended list of common contaminants. The data were searched with the following modifications: carbamidomethyl (C) as fixed modification, and oxidation (M), acetyl (protein N-term), lysine di-glycine (K-ε-GG), phosphorylated tyrosine (T) and serine (S) and acetyl-lysine (K-Ac) as variable modifications for the respective PTMs enrichments. A maximum of 3 missed cleavages were allowed for K-Ac and K-ε-GG modifications, and 2 missed cleavages were allowed for phospho enrichment. Killifish MS data from (Di Fraia et al., 2023) were reanalysed with the same Spectronaut version using the same settings. DIA PTMs data were searched against their respective spectral library using Spectronaut Professional (18.2.33, Biognosys, Zurich, Switzerland). The library search was set to 1 % false discovery rate (FDR) at both protein and peptide levels. Data were filtered for QValue percentile of 0.2, and global imputation and true automatic normalization were applied. For DIA iNeurons data, no imputation was used and local normalization was applied. Relative quantification was performed in Spectronaut for each pairwise comparison using the replicate samples from each condition using default settings. Candidates and report tables were exported from Spectronaut for downstream analysis.

**Table 1:** MS data used for library generation on Spectronaut Software.

| ***Dataset*** | ***Data used for Library*** | ***Library size*** |
| --- | --- | --- |
| Ubiquitin- Mouse | DIA / DDA | 41949 modified peptides<br>(5950 proteins) |
| Ubiquitin- iNeurons | DIA / DDA | 69359 modified peptides<br>(7374 proteins) |
| Phosphorylation - Mouse | DIA | 40711 modified peptides<br>(3630 proteins) |
| Acetylation - Mouse | DIA / DDA | 28952 modified peptides<br>(4451 proteins) |
| iNeu Proteome | DIA | 6165 proteins |

### Data processing for TMT labeled samples

For mouse brain proteome, TMT-10plex data were processed using ProteomeDiscoverer v2.0 (Thermo Fisher). Data were searched against the relevant species-specific fasta database (Uniprot database, Swissprot entry only, release 2016_01 for Mus musculus) using Mascot v2.5.1 (Matrix Science) with the following settings: enzyme was set to trypsin, with up to 1 missed cleavage. MS1 mass tolerance was set to 10 ppm and MS2 to 0.5 Da. Carbamidomethyl cysteine was set as a fixed modification and oxidation of Methionine as variable. Other modifications included the TMT-10plex modification from the quantification method used. The quantification method was set for reporter ions quantification with HCD and MS3 (mass tolerance, 10 ppm). The false discovery rate for peptide-spectrum matches (PSMs) was set to 0.01 using Percolator (Brosch et al., 2009). Reporter ion intensity values for the PSMs were exported and processed with procedures written in R (v. 3.4.1), as described in (Heinze et al., 2018). Briefly, PSMs mapping to reverse or contaminant hits, or having a Mascot score below 15, or having reporter ion intensities below 1 3 103 in all the relevant TMT channels were discarded. TMT channels intensities from the retained PSMs were then log2 transformed, normalized and summarized into protein group quantities by taking the median value. At least two unique peptides per protein were required for the identification and only those peptides with one missing value across all 10 channels were considered for quantification. Protein differential expression was evaluated using the limma package (Ritchie et al., 2015). Differences in protein abundances were statistically determined using an unpaired Student’s t test moderated by the empirical Bayes method. P values were adjusted for multiple testing using the Benjamini-Hochberg method (FDR, denoted as ‘‘adj p’’) (Benjamini et al., 1995). Proteins with adj p < 0.05 were considered as significantly affected.

### Modified peptide abundance correction

For each enrichment, PTM report tables were exported from Spectronaut. To correct the quantities of modified peptides for underlying changes in protein abundance across the age groups compared, correction factors were calculated using the aging proteome data. For each condition and protein group, the median protein quantity was calculated and then divided by the median protein quantity in the young age group. Each modified peptide was matched by protein identifier to the correction factor table. If a modified peptide was mapped to 2 or more proteins, the correction factor was calculated using the sum of the quantity of these proteins. Further, the correction was carried out by dividing peptide quantities by the mapped correction factors, and log2 transformed. Differences in peptide quantities were statistically determined using the t-test moderated by the empirical Bayes method as implemented in the R package limma (Ritchie et al., 2015).

### GO enrichment analysis

Gene Set Enrichment Analysis (GSEA) was performed using the R package clusterProfiler (Wu et al., 2021), using the function gseGO. Briefly, protein entries were mapped to the human gene name orthologues and given in input to the function to perform the enrichment. For GO term overrepresentation analysis (ORA), the topGO R package was used.

### Alignment of Ubiquitylated sites

To align ubiquitylated sites between killifish and mouse, proteins with at least one ubiquitylated site in killifish were chosen. A local alignment was conducted using protein BLAST (v2.12.0+) (Altschul et al., 1990) with default parameters between the killifish protein sequences (Nfu_20150522, annotation nfurzeri_genebuild_v1.150922) against the mouse Uniprot proteome (release 2021_04). The top 10 matches from the BLAST search were retrieved, and each modified lysine was placed into the local alignment to determine the corresponding position in the mouse protein. A specific lysine was deemed conserved if there was a lysine at the corresponding mouse position in at least one of the top 10 hits from the BLAST alignment. A ubiquitylation site was considered conserved if the same lysine mapped from killifish to mouse was also identified as ubiquitylated in the mouse dataset. The same approach was used to map mouse and killifish ubiquitylated residues onto the human proteome (release 2021_04).

